# Membrane tension mediated mechanotransduction drives fate choice in embryonic stem cells

**DOI:** 10.1101/798959

**Authors:** Henry De Belly, Philip H. Jones, Ewa K. Paluch, Kevin J. Chalut

## Abstract

Changes in cell shape and mechanics frequently accompany cell fate transitions. Yet how mechanics affects the regulatory pathways controlling cell fate is poorly understood. To probe the interplay between shape, mechanics and fate, we used embryonic stem (ES) cells, which spread as they undergo early differentiation. We found that this spreading is regulated by a beta-catenin mediated decrease in plasma membrane tension. Strikingly, preventing the membrane tension decrease obstructs early differentiation of ES cells. We further find that blocking the decrease in membrane tension inhibits endocytosis of FGF signalling components, which direct the exit from the ES cell state. The early differentiation defects we observed can be rescued by increasing Rab5a-facilitated endocytosis. Thus, we show that a mechanically-triggered increase in endocytosis regulates fate transitions. Our findings are of fundamental importance for understanding how cell mechanics regulates biochemical signaling, and therefore cell fate.

## Main

Extensive evidence points to the existence of essential feed-backs between cell fate and cell shape (1). Changes in cell shape reflect a change in the balance of forces in a cell, and are often correlated with cell state transitions (2). Yet, the mechanisms connecting cell shape and mechanics with the signaling pathways driving fate decisions remain elusive.

An example of a fate choice that correlates with a change in cell shape is the extinguishment of naïve pluripotency in the mouse embryo (3), which can be modeled with mouse embryonic stem (ES) cells in culture (4). Notably, as cells exit the ES cell state (i.e. naïve pluripotency), they generally undergo a striking shape change, from round to spread (5). The observed cell spreading during exit from naïve pluripotency suggests an alteration in force balance, or cell mechanics, and thus a potential role for mechanical signaling in the exit from naïve pluripotency. Here, we show that the shape change observed as mouse ES cells initiate differentiation is driven by a reduction in apparent membrane tension. We linked this change in membrane tension to beta-catenin, a key pluripotency factor (6). We further show that this decrease in membrane tension enhances endocytosis, specifically of fibroblast growth factor (FGF) signaling components which, through their regulation of extracellular signal-regulated kinase (ERK) activity, are necessary for exit from naïve pluripotency.

To study the interplay between cell shape and cell fate change, we first asked if cell shape correlates with fate during the early stages of exit from naive pluripotency. ES cells were cultured on gelatin coated plates and maintained in N2B27 medium supplemented with the MEK inhibitor PD0325901, the GSK3 inhibitor CHIRON, and Leukemia inhibitory factor (culture medium known as 2i+L) (4). Exit from naive pluripotency was initiated by withdrawing the inhibitors and culturing cells in N2B27 medium alone. After 24h in N2B27 (T24), cells displayed a mixture of shapes, with most cells already flat and spread, and some still round and in colonies (Fig. 1a & Supp Fig. 1). This heterogeneity is consistent with previously described asynchrony in exit from naive pluripotency (7). To identify a potential correlation with cell fate, we fixed the cells and, in cells of different shapes, quantified the expression of Nanog, a marker of naive pluripotency, and Otx2, a transcription factor upregulated in cells that have exited naive pluripotency (7). We found that Nanog levels were significantly higher, and Otx2 significantly lower, in round cells as compared to spread cells (Fig. 1b, c), thus establishing a correlation between cell shape and exit from naive pluripotency.

**Fig. 1.**
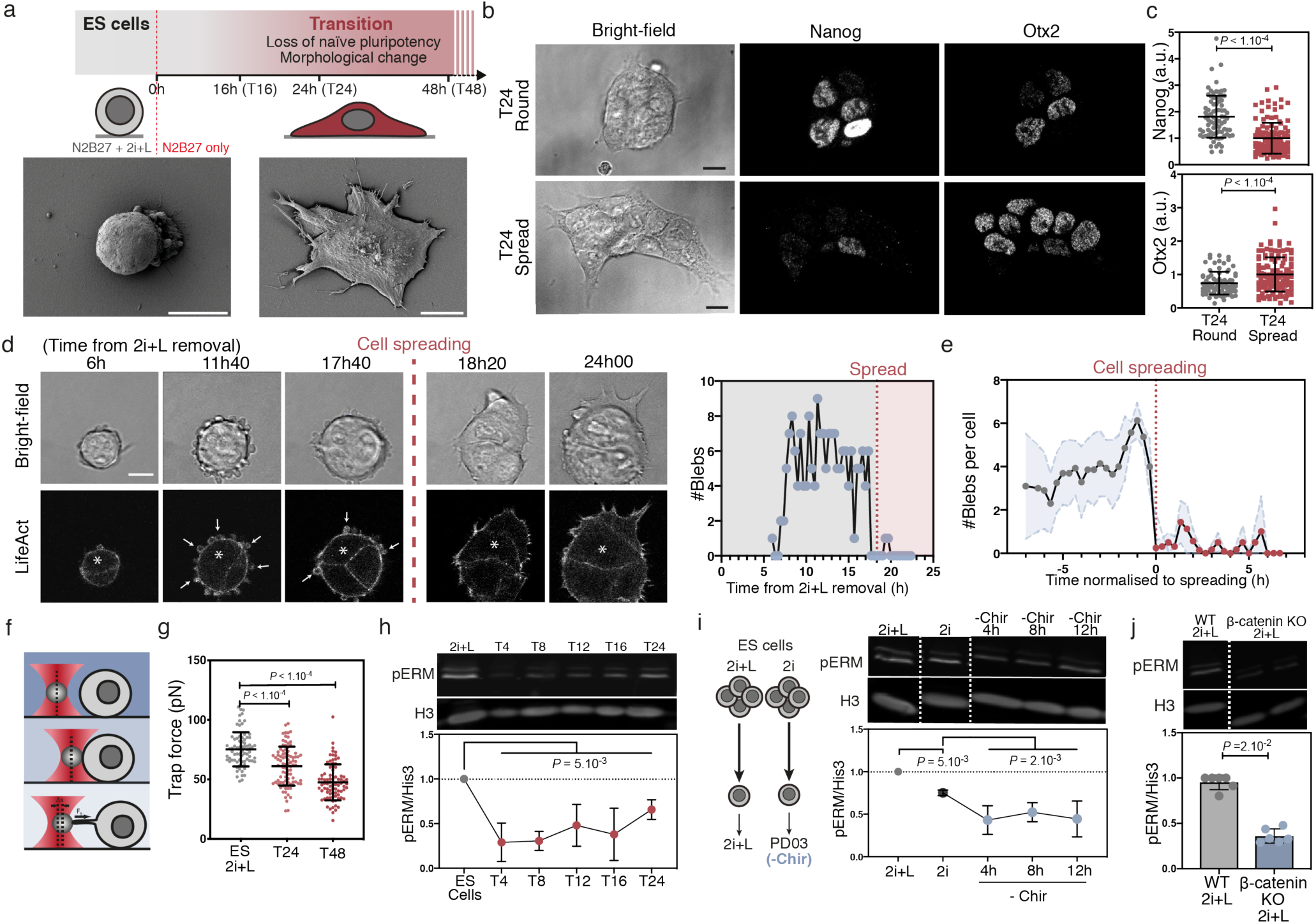
ES cells spread and show a decrease in membrane tension when exiting naive pluripotency. a, Top: schematic of experimental setup to investigate exit from naive pluripotency in ES cells; bottom: SEM images of ES and T48 cells. b, Representative single z-planes of T24 cells immunostained for Nanog and Otx2. c, Quantification of Nanog and Otx2 levels (from images as in B) in round (grey dots) and spread (red dots) T24 cells (N = 3). d, Left: time-lapse of representative ES cells expressing LifeAct-GFP and exiting naive pluripotency. White arrows: cellular blebs. Red dotted line indicates approximate time of cell spreading. Right: quantification of the number of blebs in the cell marked with a white star. e, Quantification of cell blebbing during exit from naive pluripotency. Time is normalised to the time of spreading (mean ± SD, n = 20, N=2). f, Schematic of membrane tension measurement using an optical tweezer. g, Trap force (direct readout of membrane tension) during exit from naive pluripotency (mean ± SD; 5 independent experiments). h, Representative fluorescent Western Blot for pERM and His3 in ES cells and at various timepoint during exit from naive pluripotency (see Supp Fig. 9 for uncropped Western blots) and corresponding quantification (N=4). P values are calculated using a two-way analysis of variance (ANOVA) and indicated in the figure. i, Left, schematic of the experimental setup. Right, fluorescent Western Blot and associated quantification for pERM and His3 of cells cultured in 2i+L or from 2i and replated with PD03 only for various timepoints (N=5). It is worth noting that pERM levels are lower in 2i than they are in 2i+L. P values are calculated using a two-way analysis of variance (ANOVA). j, Fluorescent Wetsern Blots and associated quantification for pERM and His3 of WT (E14) and Beta-catenin KO cells cultured in 2i+L (N=6). Graphical data represents mean ± SD. P-values: Statistical significance established by Welch’s unpaired student t-test (unless specified otherwise) and indicated in the figure. Scale bars represent 10 µm.

We next characterized the dynamics of cell spreading during exit from naive pluripotency. To do this, we captured time-lapse images of ES cells stably expressing LifeAct-GFP to visualize filamentous actin (Movie S1). We observed that, prior to spreading, the cells displayed a phase of intense blebbing (Fig. 1d, e & Supp Fig. 2a). Notably, blebbing is often a sign of low membrane-to-cortex attachment (8), a key mechanical parameter controlling the apparent tension of the plasma membrane. Apparent membrane tension (subsequently referred to as “membrane tension”) depends on the in-plane tension of the lipid bilayer and on membrane-to-cortex attachment, and is a measure of the resistance of the plasma membrane to deformations (9). As such, it is a major regulator of cell shape (10). Furthermore, in our SEM images T48 cells displayed more surface membrane folds and reservoirs compared to naive cells (Fig. 1a & Supp Fig. 1), suggesting differences in membrane tension (9). Therefore, we next asked if exit from naive pluripotency is associated with a change in membrane tension. To address this, we employed a tether pulling assay using optical tweezers (Movie S2, Fig. 1f & Supp Fig. 2b), where the force exerted on the bead by the membrane tether (the “trap force”) is a direct read-out of membrane tension (10). We found that ES cells significantly decrease their membrane tension when they exit naive pluripotency (Fig. 1g & Supp Fig. 2c). Moreover, at T24, when cell populations display mixed morphologies, round cells had a significantly higher membrane tension than blebbing and spread cells (Supp Fig. 2d, e). This is consistent with previous studies showing that cell spreading is facilitated by decreased membrane tension (11) (12). Therefore, we conclude that cell spreading during exit from naïve pluripotency occurs concomitantly with a decrease in plasma membrane tension.

**Fig. 2.**
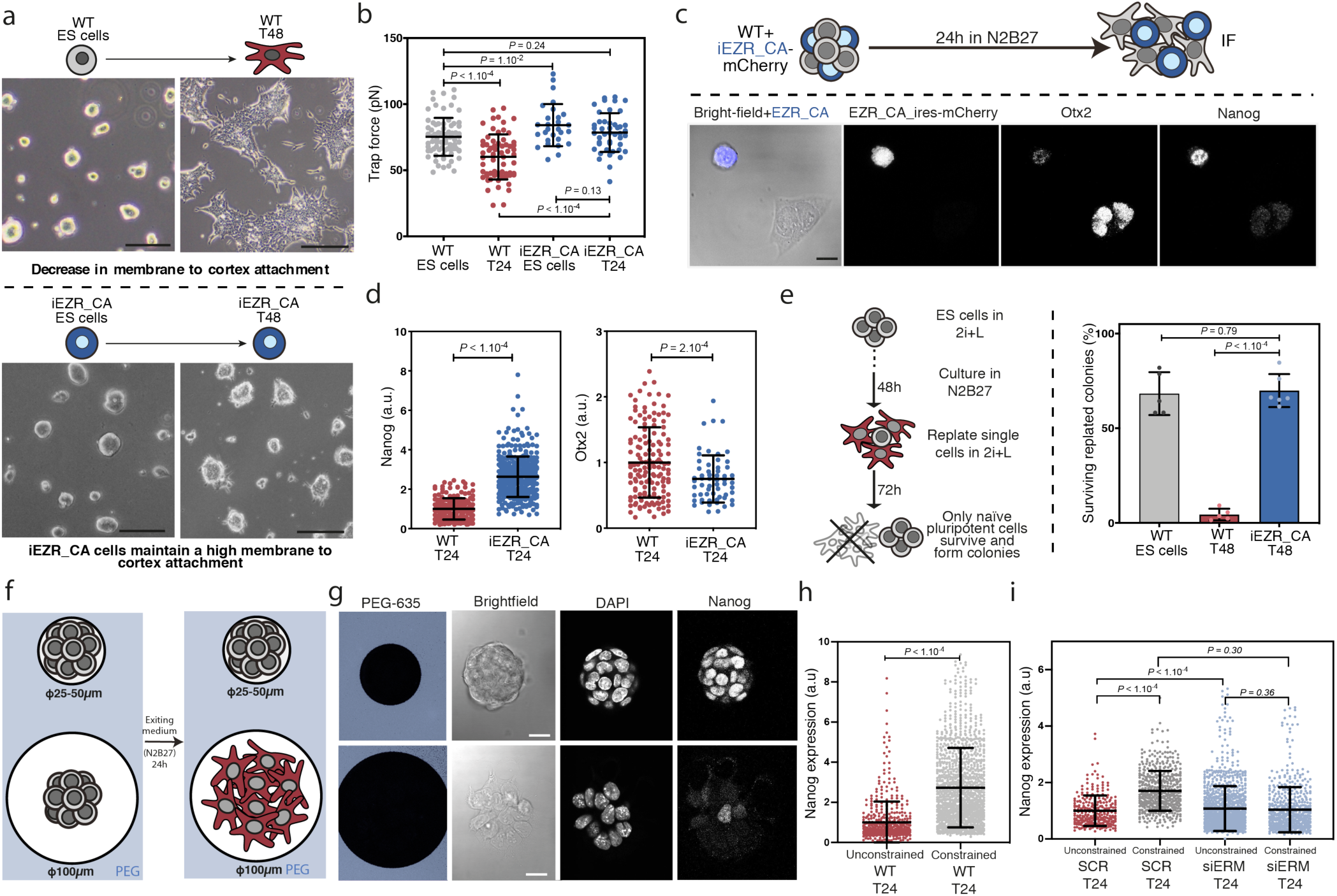
Maintaining a high membrane tension impairs exit from naive pluripotency. Representative images of WT (top) and iEZR_CA expressing (bottom) ES and T48 cells. Scale bar, 50 µm. b, Trap force, as a readout of membrane tension, for WT and iEZR_CA ES and T24 cells (N = 3; data for WT cells are from Fig. 1H). c, Representative single z-planes of a mix of WT and iEZR_CA (positive for the EZR_CA_ires_mCherry) T24 cells immunostained for Otx2 and Nanog. d, Quantification of Nanog and Otx2 expression in WT and iEZR_CA T24 cells, normalised to WT mean levels (N=3). e, Left, schematic of clonogenicity assay used to measure efficiency of exit from naive pluripotency. Right: quantification of the % of surviving replated cells in a clonogenicity assay using WT and iEZR_CA ES cells. Cells replated directly from 2i+L are used as a positive control (N=6). f, Schematic of the micropatterning assay. Cells cannot adhere on PEG regions (blue) and can only adhere on the micropatterns. g, Representative single z-plane images of fixed ES and T24 cells cultured on small (top) and large (bottom) micropatterns immunostained for Nanog. h, Quantification of Nanog expression in ES and T24 cells cultured on large (unconstrained, control) and small (constrained) micropatterns (Mann-Whitney U test was used, N = 4). i, Similar quantifications as in (h) but for cells transfected with triple siRNA against ERM or with SCR as control (Mann-Whitney U test was used, N = 4). Graphical data represents mean ± SD. P-values: Statistical significance established by Welch’s unpaired student t-test (unless specified otherwise) and indicated in the figure. Scale bars represent 10 µm.

Membrane tension has been shown to be regulated largely by linkers between the plasma membrane and the underlying actomyosin cortex, such as Ezrin-Radixin-Moesin (ERM) and Myosin I proteins (9) (13). To narrow the list of potential regulators of membrane tension, we checked the expression of genes encoding for linker proteins in ES cells and in vivo, in the pre-implantation epiblast using a published dataset (14). We found that ERM proteins, and in particular Ezrin, were up to 10 times more expressed than Myosin I proteins. We thus focused on ERM proteins, which are activated by phosphorylation (15). At the population level we found that the level of phosphorylated ERM (pERM) was sharply decreased after 2i+L removal (Fig. 1h & Supp Fig. 3a). We confirmed these results using immunostaining of T16 cells which showed that spread cells have lower levels of pERM than round cells (Supp Fig. 3 b-e). We then investigated which pluripotency-regulating signalling pathway is primarily responsible for the decrease in pERM. We first reduced the media to the minimal signalling environment necessary to maintain naive pluripotency (2i)(16). We then separately removed PD03 and Chir from 2i to study the effects of MEK/ERK activation and GSK3b activation, respectively. We found that, while PD03 removal did not lead to a pERM decrease, Chir removal resulted in a rapid and significant decrease in pERM (Fig. 1i & Supp Fig. 3f), pointing to a role for GSK3b signalling in regulating pERM levels. Given that increased GSK3b activation leads to Beta-catenin degradation (17), and that Beta-catenin is primarily localized at the plasma membrane, we asked if beta-catenin depletion would lead to a decrease in pERM. Thus, we measured pERM levels in Beta-catenin KO ES cells (6) and found that pERM is significantly lower compared to WT ES cells (Fig. 1j). These results suggest that GSK3b-driven Beta-catenin degradation mediates a decrease in pERM, which in turn controls membrane tension and cell shape during exit from naive pluripotency.

**Fig. 3.**
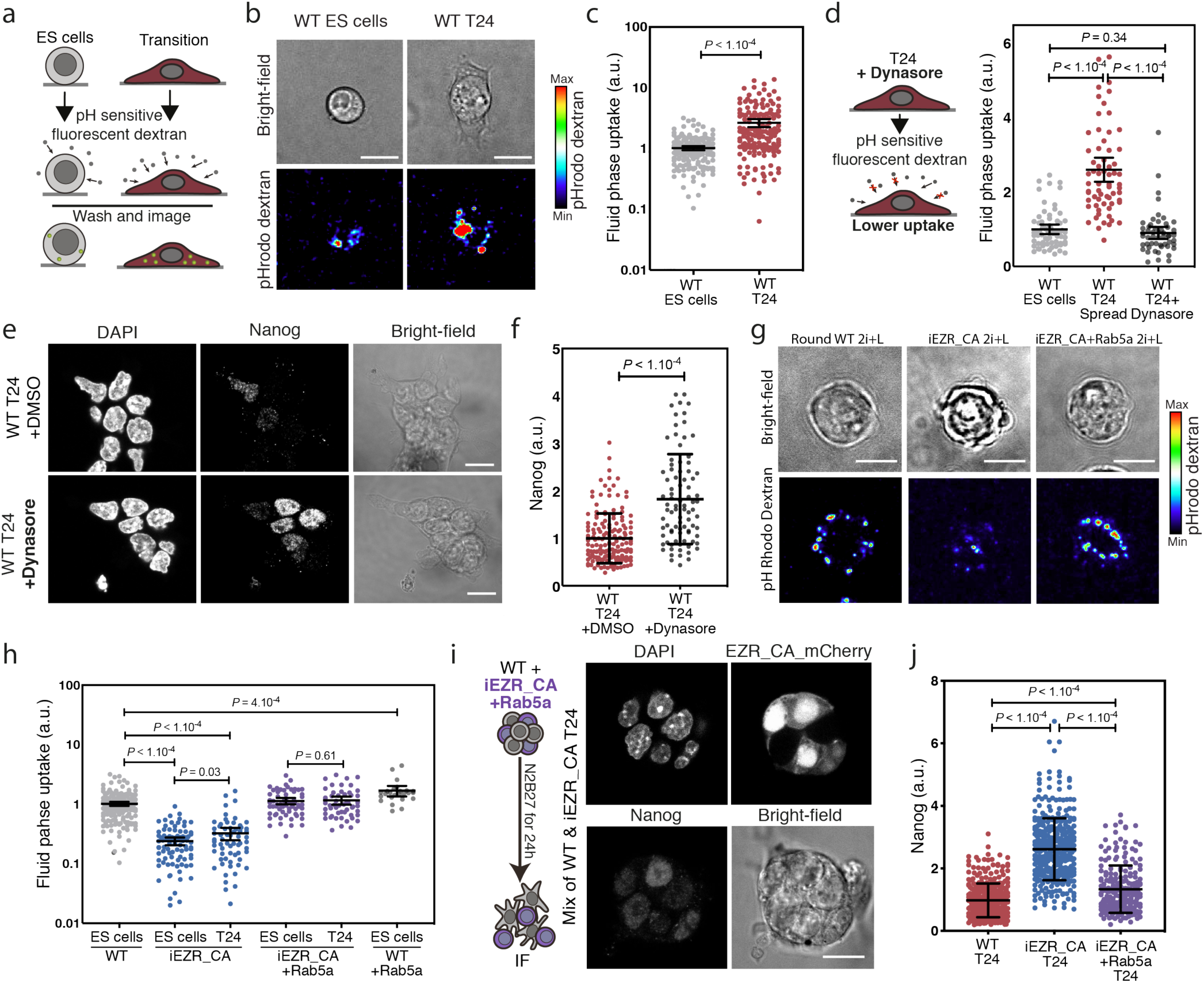
Decreasing membrane tension induces an increase in endocytosis required for exit from naive pluripotency. a, Schematic of endocytosis quantification using a fluid uptake assay with pH sensitive fluorescent dextran. b, c, Sum z-projection images of representative ES and T24 cells in assay described in a, b, and corresponding quantification (c; error bars are 95 percent Confidence IntervaIs; N=3). d, Similar assay as in a-c, for ES cells exiting naive pluripotency with either DMSO or 10 µM dynasore. (Error bars are 95% Confidence Interval; N=3). e, Representative single z-plane images of T24 cells cultured with either DMSO or 10 µM dynasore for 24h and immunostained for Nanog. f, Quantification of Nanog expression in T24 cells cultured with or without dynasore (N=3). g, h, Sum z-projection images of representative WT ES cells, iEZR_CA-ES cells and iEZR_CA-ES cells transfected with Rab5a in assay described in a and associated quantifications. Error bars: 95% Confidence Intervals (N=3). i, j, Single z-plane images of representative mixed populations of WT and iEZR_CA (positive in the mCherry channel) T24 cells, transfected with Rab5a and immunostained for Nanog and corresponding quantifications (N=3). Non transfected iEZR_CA data are from Fig. 2D. Graphical data represents mean ± SD. P-values: Statistical significance established by Welch’s unpaired student t-test (unless specified otherwise and indicated in the figure. Scale bars represent 10 µm.

We then asked if the membrane tension decrease plays a role in controlling exit from naive pluripotency. We used a Doxycycline-inducible ES cell line expressing a phospho-mimetic, constitutively active form of Ezrin (EZR_CA). After induction (iEZR_CA), cells showed high EZR_CA expression and no growth defects (Supp Fig. 4a, b).After withdrawal of the naive factors (2i+L), iEZR_CA cells did not spread (Fig. 2a), even at T48, and they maintained high membrane tension similar to ES cells (Fig. 2b). We also found that, compared to controls, iEZR_CA cells at T24 maintained significantly higher levels of Nanog and failed to efficiently upregulate Otx2 (Fig. 2c, d & Supp Fig. 4c, d). A clonogenicity assay indicated that unlike control T48 wild type cells, iEZR_CA cells placed back in 2i+L after 48 hours in N2B27, were able to survive and form naive colonies with the same efficiency as naive ES cells (Fig. 2e, see Materials and Methods for details). These results indicates that the iEZR_CA cells are significantly inhibited in their ability to exit naive pluripotency. Furthermore, treating cells with methyl-β-cyclodextrin, a cholesterol-sequestering compound, which increases membrane tension (18) (Supp Fig. 4e, f), delayed exit from naive pluripotency (Supp Fig. 4g). Conversely, treating cells with NSC 668394, an ezrin inhibitor which decreases membrane tension19 (Supp Fig. 4e, f), caused cells to exit naive pluripotency more efficiently than control cells (Supp Fig. 4g). Taken together, our data suggest that a pERM-controlled decrease in plasma membrane tension is essential for cell spreading and for exit from naive pluripotency.

**Fig. 4.**
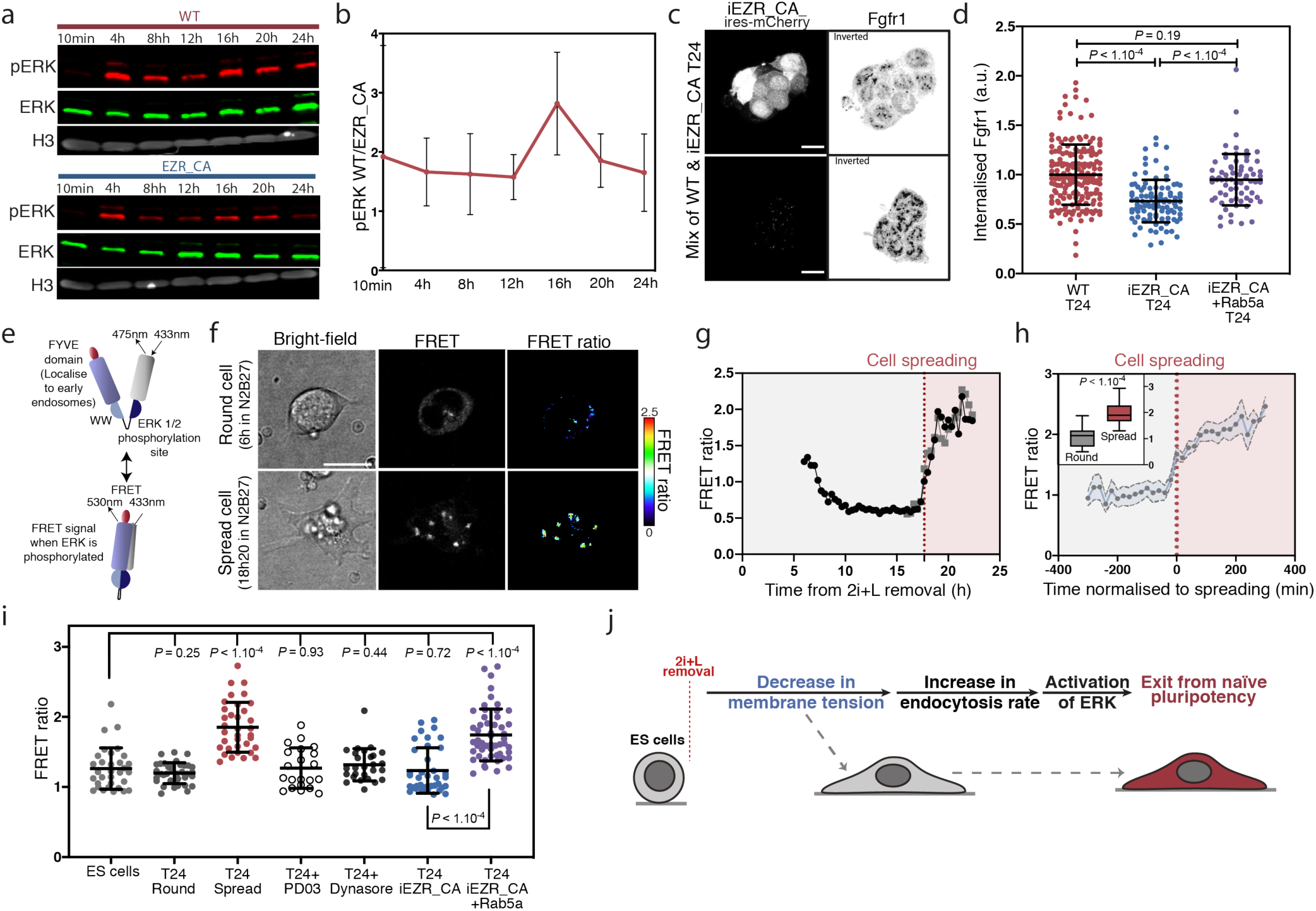
Decrease in membrane tension and increased endocytosis promote ERK activation during exit from naive pluripotency. a, Fluorescent Western blot for ERK, pERK and Histone3 in ES cells at different timepoints during exit from naive pluripotency (see Supp Fig. 9 for uncropped Western blots). b, Time-course of the ratio of pERK levels (normalised to corresponding Histone3 levels) in WT and iEZR_CA cells during exit from naive pluripotency (N=6). c, Representative single z-plane images of a mixed population of WT and iERC_CA (positive in the mCherry channel) T24 cells, immunostained for Fgfr1. d, Quantification of average intracellular Fgfr1 levels in T24 cells across different conditions (N=3). e, Schematic of the FRET ERK sensor used to measure ERK activation at early endosomes 26. f, Time-lapse of representative cell expressing the FRET ERK sensor exiting pluripotency at 6 h (top, the cell is still round) and 18h20 (bottom, the cell is spread). g, Timecourse of the FRET ratio for the cell shown in f. h, Quantification of the FRET ratio in ES cells exiting naive pluripotency, time is normalised to the time of spreading (means ± SEM, n=10, N=3). Inset: FRET ratio for round and spread cells in the same dataset. i, Mean FRET ratio for fixed ES and T24 cells in different conditions. Dynasore was used at 10µM. PD03 was used at 1µM. (N=3). j, Schematic of the proposed mechanism for how membrane tension regulates signalling during exit from naive pluripotency. Graphical data represents mean ± SD. P-values: Statistical significance established by Welch’s unpaired student t-test (unless specified otherwise) and indicated in the figure. Scale bars represent 10 µm.

To elucidate the relative importance of cell spreading and membrane tension, we used micropatterning to control the extent of cell spreading with disks of different diameters (small: 25-50 µm or large: 100 µm). We seeded ES cells at equal densities onto both small and large micropatterns in N2B27 only. We found that in contrast to cells plated on large patterns or non-patterned surfaces, spatially-constrained T24 cells on small patterns did not change shape and maintained a rounded morphology similar to ES cells (Fig. 2f, g). Spatially constrained cells at T24 maintained high levels of Nanog suggesting defects in exiting naive pluripotency (Fig. 2h Supp Fig. 5a). Notably spatially constrained cells also maintained a high membrane tension (Supp Fig. 5b). Strikingly, the exit from naive pluripotency on small micropatterns, where cells cannot spread, was rescued by knockdown of ERM (Fig. 2i & Supp Fig. 5c-f) or knockdown of Myosin I family proteins (Supp Fig. 5g-i), treatments previously shown to reduce membrane tension (19) (20). Together, these experiments point to the existence of positive feedback loops between membrane tension decrease and cell spreading, and suggest that it is the decrease in membrane tension, rather than cell spreading, is required for efficient exit from naive pluripotency.

To further test if membrane tension regulates exit from naive pluripotency directly, or as a result of cell spreading, we plated ES cells on laminin, where they display slightly spread morphologies even in 2i+L (Supp Fig. 5j). Notably, despite the fact that for cells on laminin exit from pluripotency was associated with a much less pronounced spreading increase than for cells on gelatin, we still observed a drop in membrane tension during exit from pluripotency (Supp Fig. 5k). This further indicates that it is the decrease in membrane tension, and not the spreading directly, that is needed for efficient dismantling of naive pluripotency. Taken altogether, our results strongly implicate a role for Ezrin-mediated drop in membrane tension in exit from naive pluripotency.

We then asked how decreased membrane tension facilitates exit from the ES cell state. We speculated that the facilitation may be through endocytosis, which is a major regulator of signalling events (21)(22) and has been shown to be regulated by membrane tension (23) (24). First, we assessed global endocytosis levels using a fluid phase uptake assay, in which we cultured cells in presence of either fluorescent dextran or a pH-sensitive fluorescent dextran (Fig. 3a). Using this assay, we found that T24 cells displayed a significant increase in endocytosis levels once spread (Fig. 3b, c & Supp Fig. 6a-d). Blocking dynamin-mediated endocytosis with dynasore (25) (Fig. 3d & Supp Fig. 6e), resulted in cells that maintained a higher level of Nanog protein expression (Fig. 3e, f) and colony-forming ability in 2i+L (Supp Fig. 6f) compared to controls. These observations indicate that endocytosis plays an important role in exit from naive pluripotency.

We next examined the link between the observed differentiation-induced drop in membrane tension and endocytosis. First, we showed that endocytosis was considerably reduced in cells expressing iEZR_CA (Fig. 3g, h), suggesting that high membrane tension antagonizes endocytosis, consistent with previous findings in other systems(23) (24). To test whether enhanced endocytosis could overcome the block in differentiation imposed by EZR_CA, we transfected iEZR_CA cells with Rab5a, a master regulator of early endosomes (26). Elevation of Rab5a levels has been shown to promote clathrin-independent endocytosis (26) (27). Indeed, we found that overexpressing Rab5a increased endocytosis levels in iEZR_CA cells compared to non-transfected controls, both in ES and T24 cells (Fig. 3g), without lowering membrane tension (Supp Fig. 7g). Notably, Rab5a expression rescued the exit phenotype in iEZR_CA cells, restoring Nanog downregulation (Fig. 3i, j) and decreasing the number of colonies surviving in a clonogenicity assay, indicative of more efficient exit from naive pluripotency (Supp Fig. 6h). Furthermore, transfection with Rab5a also enabled downregulation of Nanog protein in spatially-constrained T24 cells (Supp Fig. 6i, j). Altogether, these results show that increasing endocytosis levels via Rab5a overexpression allows ES cells to exit naive pluripotency notwithstanding a high membrane tension.

We next asked if a specific signalling pathway related to regulation of naive pluripotency was affected by the changes in cell mechanics and endocytosis we observed. The most obvious candidate was FGF/ERK, as it is key for exit from naive pluripotenc (28), and has been shown to be regulated by endocytosis in cancer cells (22) (26). We thus quantified pERK levels during exit from naive pluripotency, and observed that at all time points ERK activity was highly attenuated in the iEZR_CA cells (Fig. 4a, b & Supp Fig. 7a, b). We confirmed a reduction in MAPK/ERK signalling in iEZR_CA cells by qPCR on known transcriptional targets (Supp Fig. 7c), strongly suggesting that maintaining a high membrane tension during exit from naive pluripotency prevents ERK activation.

Internalization of receptors by endocytosis has been shown to influence the levels pERK activity (26) (29). Thus, we measured internalization of FGFR1, which is upstream of pERK in ES cells (30), in iEZR_CA and control cells using immunofluorescence. We found that iEZR_CA cells had less internalized FGFR1 compared to controls (Fig. 4c, d), despite expressing similar protein levels (Supp Fig. 7d, e). We were able to increase the levels of internalized FGFR1 in iEZR_CA by over-expressing Rab5a (Fig. 4c, d). We then specifically probed pERK activity at endosomes using a recently described FRET sensor reporting on ERK activity at early endosomes (Fig. 4e) (26). Using live imaging, we found that pERK at endosomes sharply increased as cells spread (Supp Fig. 4f-h and Supp Fig. 8, Movie S3). We confirmed this with measurements in fixed cells, finding that spread T24 cells showed a higher FRET ratio compared to round T24 cells. Notably, we observed a lower FRET ratio in T24 cells where endocytosis rates were reduced chemically with dynasore or by increasing membrane tension via iEZR_CA expression (Fig. 4i). Furthermore, increasing endocytosis independently of membrane tension, by expression of Rab5a in iEZR_CA cells, resulted in a FRET ratio similar to spread control cells at T24 (Fig. 4i). Taken all together, our results indicate that ERK activity, which helps disassemble naive pluripotency, is induced by membrane tension-regulated endocytosis in cells exiting pluripotency (Fig. 4j).

We have found that a decrease in plasma membrane mechanics upon exit from pluripotency activate FGF/ERK via enhanced endocytosis of FGF receptors. The fact that the decrease in membrane tension is regulated by degradation of Beta-catenin, suggests a previously unknown synergy between the two essential pluripotency regulators Beta-catenin and FGF/ERK. Our finding suggests that activation of ERK signalling through FGF will not be as efficient if there are high levels of activated beta-catenin.

Taken together, our work unveils a mechanism directly connecting a change in cell mechanics to the activation of a biochemical signalling pathway. We anticipate that the interplay between membrane tension and endocytosis-mediated enhanced signalling could be an important regulator of cell fate in various developmental and pathological contexts.

## ACKNOWLEDGEMENTS

We thank Carla Mulas for comments on the manuscript, Giorgio Scita for advice on endocytosis, Ortrud Warlick for help with setting up the optical tweezer, the MRC LMCB light microscopy facility for technical support, Anan Ragab and Mark Turmaine for their help with SEM, members of the Paluch lab (in particular Agathe Chaigne and Murielle Serres) and of the Chalut lab (in particular Ayaka Yanagida) for technical help and advice at different stages of this project.

## Supplementary Materials

### Methods

#### Cell culture, transfections and exit from naïve pluripotency

Mouse embryonic stem cells were cultured on Falcon flasks coated with 0.1% gelatine. On occasion, laminin (Sigma-Aldrich #11243217001) coated dishes were used instead of gelatine. ES cells used were E14TG2a. Cells were cultured in N2B27+2i+LIF (2i+L) (4) at 37°C with 7% CO_2_. Cells were passaged every other day using Accutase (Sigma-Aldrich, #A6964) and regularly tested for mycoplasma. Culture media was changed every 24h. The culture media was made using DMEM/F-12, 1:1 mixture (Sigma-Aldrich, #D6421-6), Neurobasal medium (Life technologies #21103-049), 2.2 mM L-Glutamin, B27 (Life technologies #12587010), 3 µM Chiron (Cambridge Bioscience #CAY13122), 1 µM PD0325901 (Sigma-Aldrich #PZ0162), 20 ng/ml of LIF (Merck Millipore # ESG1107), 50 mM β-Mercapto-ethanol, 12.5 ng.mL^-1^ Insulin zinc (Sigma-Aldrich #I9278) and home-made N2. The 200 X home-made N2 was made using 0.791 mg.mL^-1^ Apotransferrin (Sigma-Aldrich #T1147), 1.688 mg.mL^-1^ Putrescine (Sigma-Aldrich #P5780), 3 μM Sodium Selenite (Sigma-Aldrich #S5261), 2.08 µg.mL^-1^ Progesterone (Sigma-Aldrich #P8783), 8.8% BSA. Exit from naïve pluripotency was triggered by passaging ∼500 000 cells and seeding them in N2B27 only (N2B27) on T25 gelatine coated flasks.

Transfections of plasmids were performed using 5 μg of plasmid and 3.5 μL of Lipofectamine 2000 (ThermoFischer Scientific #11668019), incubated in 250 μL OptiMEM for 5 minutes, then mixed and incubated at room temperature for 30 minutes, and added to cells passaged onto Ibidi dishes with polymer coverslip bottom for observations(IBIDI Scientific, 81156).

siRNA transfections were performed using 30 μM of siRNA and 5 μL of Lipofectamine RNAiMAX (ThermoFischer Scientific #13778075) incubated in 250 μL OptiMEM for 5 minutes, then mixed and incubated at room temperature for 30 minutes and added to cells passaged onto micropatterns.

#### Cell lines

ES-E14TG2a (referred as WT) directly derived from mouse embryos were a kind gift from the Austin Smith’s laboratory (Cambridge Stem Cell Institute). The LifeAct GFP cell line is from (31). These cells have been generated and kindly given to us by Ian Rosewell (Francis Crick Institute, under Holger Gerhardts project license). Dox-inducible IRES mCherry Ezrin_CA ES cells were kindly given to us by Ayaka Yanagida from the Chalut and Nichols’s lab. The EZR_CA was engineered using a PiggyBac system with E14 ES cells. The Beta-Catenin Knockout cell line was kindly given to us by the lab of Austin Smith and has been described in (6).

#### Tether pulling and trap force measurements

Trap force measurements were performed using a home-built optical tweezer using a 4W 1064 nm Laser Quantum Ventus with a 100x oil immersion objective (NA 1.30, CFI Plan Fluor DLL, Nikon) on an inverted microscope (Nikon Eclipse TE2000-U) equipped with a motorized stage (PRIOR Proscan). The optical tweezer was calibrated following (32). Measurements were performed using concanavalin-A coated (50 µg/ml) carboxyl latex beads (1.9µm diameter, Thermo Fisher C37278). Beads were incubated on a shaker with concanavalin-A for one hour prior to the experiment. Bead position was recorded every 90 milliseconds in bright field prior and during tether formation. The trap force was calculated based on the calibration of the trap and the bead position using a home-made Fiji (33) plugin (HDB) (typical values for the trap stiffness were k∼0.130 pN/nm.)

#### RNA extraction and qPCR

Cells were seeded in 6 well dishes in different conditions (N2B27 or 2i+L) and cultured for different periods of time, as indicated. RNA was extracted using a Qiagen (205310) kit. Extracted RNA cDNA was generated from the RNA according to kit’s instructions (Thermo Fischer Scientific #4368814). For the RT-qPCR, pre-designed primers (see table of primers used), and the SYBR Green Master Mix (Qiagen; 204141) were combined according to the kit’s instructions. qPCR was performed on Bio-Rad CFX qPCR.

#### Western blots

Cells were plated into 6-well plates in different conditions (with or without 2i+L), as indicated. Proteins were extracted on ice using Laemmli buffer (Bio-Rad #1610747). Samples were put at 95° C for 5 min and then sonicated on ice. Isolated whole protein content was measured using the Thermo Scientific Pierce 660nm Protein assay (#22660) using a BSA gradient kit (Bio-Rad; 500-0206) and 20 µg of protein extract was loaded onto either NuPAGE 4-12% Bis-Tris protein gels (Thermo Fischer Scientific #NP0321BOX) or Novex WedgeWell 14% Tris-Glycine Mini Gels (Thermo Fischer Scientific #XP00140BOX) depending on the size of the protein of interest. Gels were run at 125 V for 2 h. Proteins were transferred for 70 minutes at 70 V to a nitrocellulose membrane (Thermo Fischer Scientific #88024). Membranes were blocked for 90 min using Odyssey blocking buffer (Licor #927-50003). Primary antibodies were added at the proper dilution (see table of antibodies) in either TBS-T + 5% milk or 5% BSA and incubated overnight at 4 degrees. Membranes were washed three times for 15 min in TBS with TWEEN. Infrared species-appropriate secondary antibodies were then added (see table of antibodies) for one hour. Membranes were washed three times with PBS-T for 15 min and imaged on the Licor Odyssey scanner. Bands were quantified using LI-COR Biosciences Image Studio Lite.

#### Immunofluorescence

Cells were fixed in IBIDI dishes (IBIDI Scientific, 81156) with 4% formaldehyde. Cells were permeabilized during 10 min in PBS+0.1% Triton-X. Blocking was then done using with 2% FBS 2% BSA in PBS with 0.1% Triton for 45 min. Cells were then incubated for 1h30min with the primary antibody diluted at the appropriate concentration in the same buffer as used for blocking. Cells were then washed 3 times 5 min with PBS+0.1% Triton. Secondary antibodies were added for 1 h diluted at 1:800 in the same buffer as used for blocking and primary antibody incubation. Cells were washed with PBS + 0.1% Triton 3 times for 5 min. Finally, cells were incubated for 5 min with PBS+DAPI before a final rinse in PBS.

#### Quantifications

Live imaging was performed on an inverted confocal microscope (Olympus FV1200), at 37°C with 5% CO_2_, taking z-stacks 10μm in total height with a 1μm step using a 63x objective and using low laser power. Fix imaging was performed on an inverted confocal microscope (Leica TCS SP5) taking z-stacks 20μm in total height with a 1μm step using a 63x objective. All image analysis were performed using Fiji (33).

For Nanog and Otx2 levels quantifications, average fluorescence was measured on the midplane of the nucleus. DAPI was used as a marker to locate the nuclei.

For pERM and ERM levels quantifications in Fig S3, total fluorescence was measured in each cell following a sum z projection. Signal was corrected for background by taking a ROI of the background of a similar size as of the measured cells. The background signal was then subtracted from the signal from cell. Data were normalised to round cells. Alexa Fluor 568 Phalloidin (ThermoFischer Scientific # A12380) signal was used to determine cell boundaries and to classify cells according to shape based on visual inspection.

For Fgfr1 quantifications, average fluorescence was measured in each cell using CellMask Deep Red (Invitrogen # C10046) as a marker to locate individual cells. Average intensity was then manually measured in the midplane of the cells by taking a ROI following the cell boundary.

For #blebs in Fig. 1, the number of blebs was manually counted at every time point (every 20 minutes) across a z-stack using LifeAct and brightfield as guides to identify individual blebs (note that only actin filled blebs could be observed with this method).

#### Micropatterning

Glass coverslips (Marienfekd GmBH) were sonicated in 1 M of HCl for 5-15 minutes, washed and plasma-treated for 30 seconds for glass passivation. Then 0.1 mg/mL of PLL-g-PEG (PLL(20)-g[3.5]-PEG(2)/Atto663 from SuSoS) solution was added to the coverslips for 30 min at room temperature. Chrome MASK from Delta Mask was used. The mask was illuminated with UV light for 5 min on the chrome side, then the PEG-coverslips were added on the chrome side and illuminated for 6 min to burn the PEG in the shape of the desired pattern (circles in our case). Coverslips were then dried and stored. Coverslips were rehydrated before use.

For immunofluorescence assays, we seeded the cells onto the patterns (coated with laminin, see above for details) in 2i+L at a density allowing for the formation of small colonies, occupying the whole available space on the small circles while still having free space on the bigger circles. Cells were then washed three times in PBS and placed in N2B27 only without 2i+L. After 24h in N2B27, cells were fixed and immunostained.

#### Fluid phase uptake assay

Cells were seeded on IBIDI dishes (IBIDI Scientific, 81156) a few hours before imaging. Cells were then incubated for 10 min in N2B27 or 2i+L with 10 μg of either pH-Rhodo Dextran (pHrodo™ Red Dextran, 10,000 MW, for Endocytosis, Thermofischer Scientific) or Alexa Fluor Dextran (Dextran, Alexa Fluor™ 647; 10,000 MW, Thermofischer Scientific) (34). Cells were then rinsed immediately prior to imaging. Each dish was imaged for 30 min.

For quantifications with either pH Rhodo or Regular Dextran, total intensity inside the cell was measured with Fiji by using a sum z projection. The signal was corrected by taking a ROI of the background of a similar size as of the measured cells. The background signal was then subtracted from the signal from cell. Only single cells or groups of cells with clear boundaries were quantified. CellMask Deep Red (Invitrogen # C10046) was used to identify individual cell boundaries. Experiments were always performed side by side with cells in 2i+LIF as control. Data are normalised to cells in 2i+L.

#### FRET analysis

Imaging was performed on an inverted confocal microscope (Olympus FV1200), at 37°C with 5% CO_2_ for live imaging, taking z-stacks 10μm in total height with a 1μm step using a 63x objective. For FRET quantifications, we used a home-written (HDB) custom Macro in Fiji [5] FRET and donor channel signals were corrected for background, vesicles were then semi-automatically identified and average fluorescence was quantified in each vesicle across all the channels. Small Z maximum projections (over 3 z slices of 1μm) were quantified in order to ensure capture entire endocytotic vesicles. We then calculated the ratio of values obtained in donor and FRET channels to obtain FRET ratio following (35) (26) methods.

#### Clonogenicity assay

To test for speed and efficiency of exit from pluripotency, replating assays were performed. For each experimental condition, ∼400 000 cells were plated onto 6-well dishes coated with 0.1% gelatin, in N2B27 to induce exit from naïve pluripotency. After 48h in N2B27, the cells were resuspended and counted. A specific number of cells (typically 400) was then replated in N2B27+2i+L onto a 12-well plate. After 4 days, the number of colonies was manually counted; as only naïve cells survive in 2i+L, this assay quantifies the efficiency of pluripotency exit. As a positive control, cells cultured in 2i+L were replated in 2i+L. Occasionally replated cells were tested for naïve pluripotency using a Blue-colour AP staining kit to add another control that replated cells were indeed naïve pluripotent (Cambridge Bioscience #AP100B-1).

#### Statistical analysis

For all statistical analysis, PRISM 7 (Graphpad software, Inc) was used. Unless specified all test in figures are unpaired student t-tests with Welch’s correction. D’agostino and pearson test was used to test for normality.

#### Scanning Electron Microscopy

Samples were fixed with glutaraldehyde 1%. Preparation for scanning electron microscopy was performed as in (<u>36</u>) without membrane extraction. Jeol 7401 - high resolution Field Emission Scanning Electron Microscope was used for imaging.

#### Table of antibodies

| Antibody | Cat No | Company | Dilution |
| --- | --- | --- | --- |
| phospho ERK | 4370 S | Cell Signaling | 1:1000 (WB) |
| ERK | 9107 S | Cell Signaling | 1:1000 (WB) |
| Nanog | eBioMLC-51 | eBioscience | 1:200 (IF) |
| Otx2 | AF1979 | R&D Systems | 1:200 (IF) |
| ERM | 3142S | Cell Signalling | 1:2000 (WB)<br>1:200 (IF) |
| Phospho ERM | 3149S | Cell Signaling | 1:2000 (WB)<br>1:200 (IF) |
| GAPDH | ab8245 | Abcam | 1:5000 (WB) |
| His3 | Ab1701 | Abcam | 1:5000 (WB) |
| Fgfr1 | ab10646 | Abcam | 1:2000 (WB)<br>1:200 (IF) |

#### Plasmids

Rab5a and FRET plasmids were provided by Giorgio Scita’s lab (26)

Ezr_GFP and EZR_CA_GFP were provided by Guillaume Charras (15)

#### Primers

Dusp6:

5’ - GCTGATACCTGCCAAGCAAT– 3’

5’ – TCCTATCTGGATCACTGGAG – 3’

Spry4:

5’ – TGCACCATGACTGAGCTG – 3’

5’ – AGCGATGCTTGTGACTCTG – 3’

Spry2:

5’ – CATCAGGTCTTGGCAGTGT– 3’

5’ - CATCGCTGGAAGAAGAGGAT– 3’

Egr1:

5’ – GATAACTCGTCTCCACCATCG – 3’

5’ – AGCGCCTTCAATCCTCAAG - 3’

#### Table of reagents and siRNA

| Reagents | Cat No | Company | Concentration used |
| --- | --- | --- | --- |
| Hoechst | 34580 | ThermoFischer | 200 ng/ml |
| DAPI | 9542 | Sigma-Aldrich | 200 µg/ml |
| Dynasore | D7693 | Sigma-Aldrich | 10 µM |
| NSC 23766 | 2161 | Tocris | 25 µM |
| Methyl-B-Cyclodextrin | C4555 | Sigma-Aldrich | 1 mM |
| Dextran, Alexa Fluor™ 647,<br>10,000 MW | D22914 | thermofischer | 10 µg |
| pHrodo™ Red Dextran,<br>10,000 MW | P10361 | thermofischer | 10 µg |
| SMARTpool: ON-TARGETplus Ezr siRNA | L-046568-01 | Dharmacon | 30 nM |
| SMARTpool: ON-TARGETplus Rdx siRNA | L-047230-01 | Dharmacon | 30 nM |
| SMARTpool: ON-TARGETplus Msn siRNA | L-044428-01 | Dharmacon | 30 nM |
| SMARTpool: ON-TARGETplus MYO1C<br>siRNA | L-015121-00 | Dharmacon | 30 nM |
| SMARTpool: ON-TARGETplus Myo1b siRNA | L-045103-01 | Dharmacon | 30 nM |
| siGENOME Non-Targeting<br>Control siRNA #3 | D-001210-03 | Dharmacon | 10 nM |
| siGENOME Non-Targeting<br>Control siRNA #5 | D-001210-05 | Dharmacon | 10 nM |
| siGENOME Non-Targeting<br>Control siRNA #2 | D-001210-02 | Dharmacon | 10 nM |

---

**Movie S1.**

Time-lapse (confocal) of LifeAct ES cells exiting naïve pluripotency. Movie starts at 6 h after media change to N2B27. One frame is shown every 20 min.

**Movie S2.**

Time-lapse (brightfield) of a ES cell where a membrane tube is pulled with an optical tweezer. One frame is shown every second. A red target has been added to visualize the center of the laser trap.

**Movie S3.**

Time-lapse (confocal) of ES cells exiting naïve pluripotency and transfected with FRET sensor for ERK activity. Movie starts after 6 h in N2B27. One frame is shown every 20 minutes. A single z-slice is shown.

## Funding

This work was supported by the European Union’s Horizon 2020 research and innovation programme under the Marie Sklodowska-Curie grant agreement No *641639* (ITN Biopol, HdB and EKP), the Medical Research Council UK (MRC programme award MC_UU_12018/5, HdB and EKP), the Human Frontier Science Program (Young Investigator Grant RGY 66/2013 to EKP), the Leverhulme Trust (Prize in Biological Sciences to EKP), a core support grant from the Wellcome Trust and Medical Research Council to the Wellcome Trust - Medical Research Council Cambridge Stem Cell Institute (KJC). KJC is a Royal Society University Research Fellow.

## Data and materials availability

All data is presented in the main text or the supplementary materials.

---

**Supp Fig. 1:**
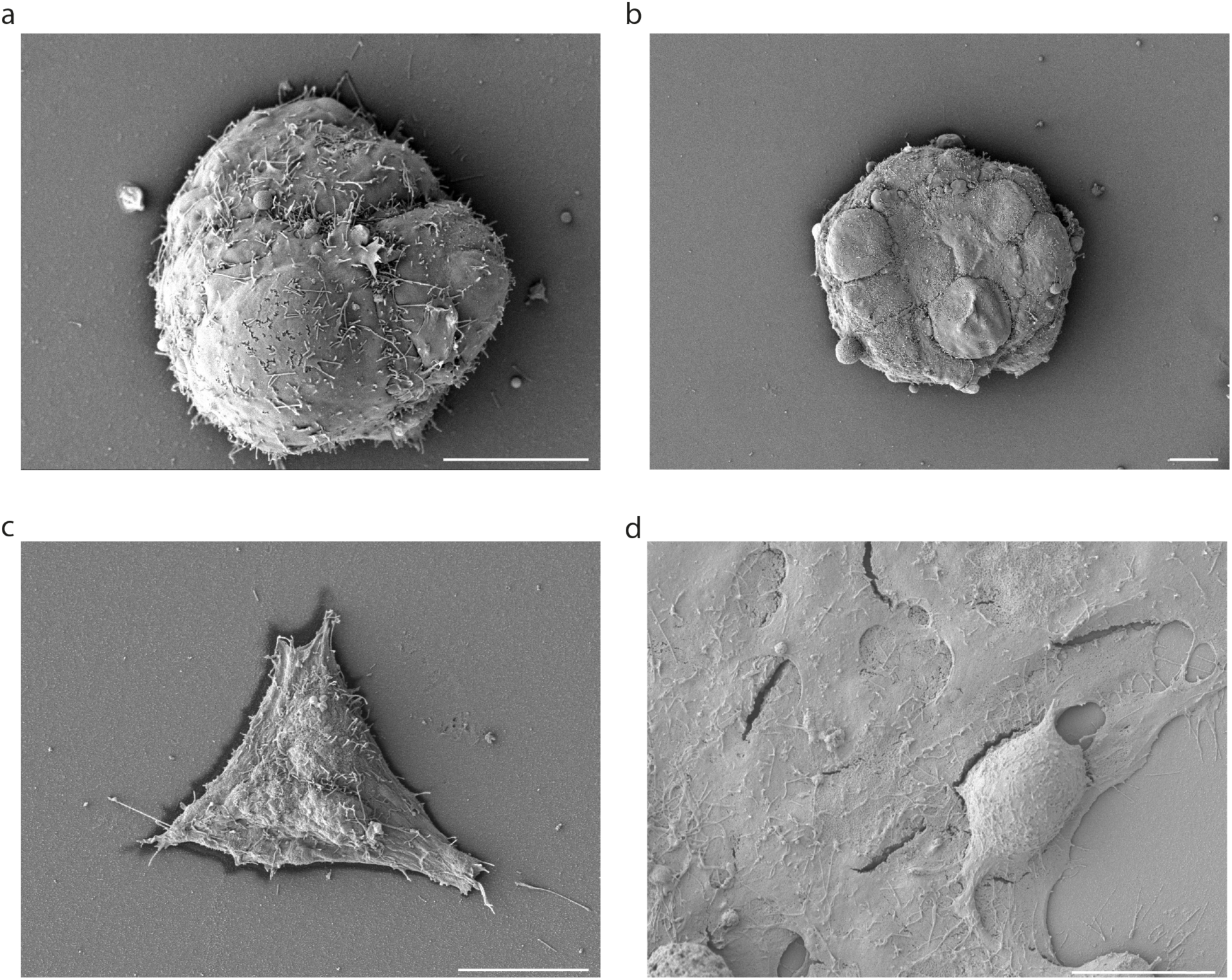
SEM images of naïve and transition ES cells. Representative SEM images of ES cells. a, b, colonies of ES cells. c, a single T48 cell. d, and colonies of T48 cells. Scale bars, 10 μm.

**Supp Fig. 2:**
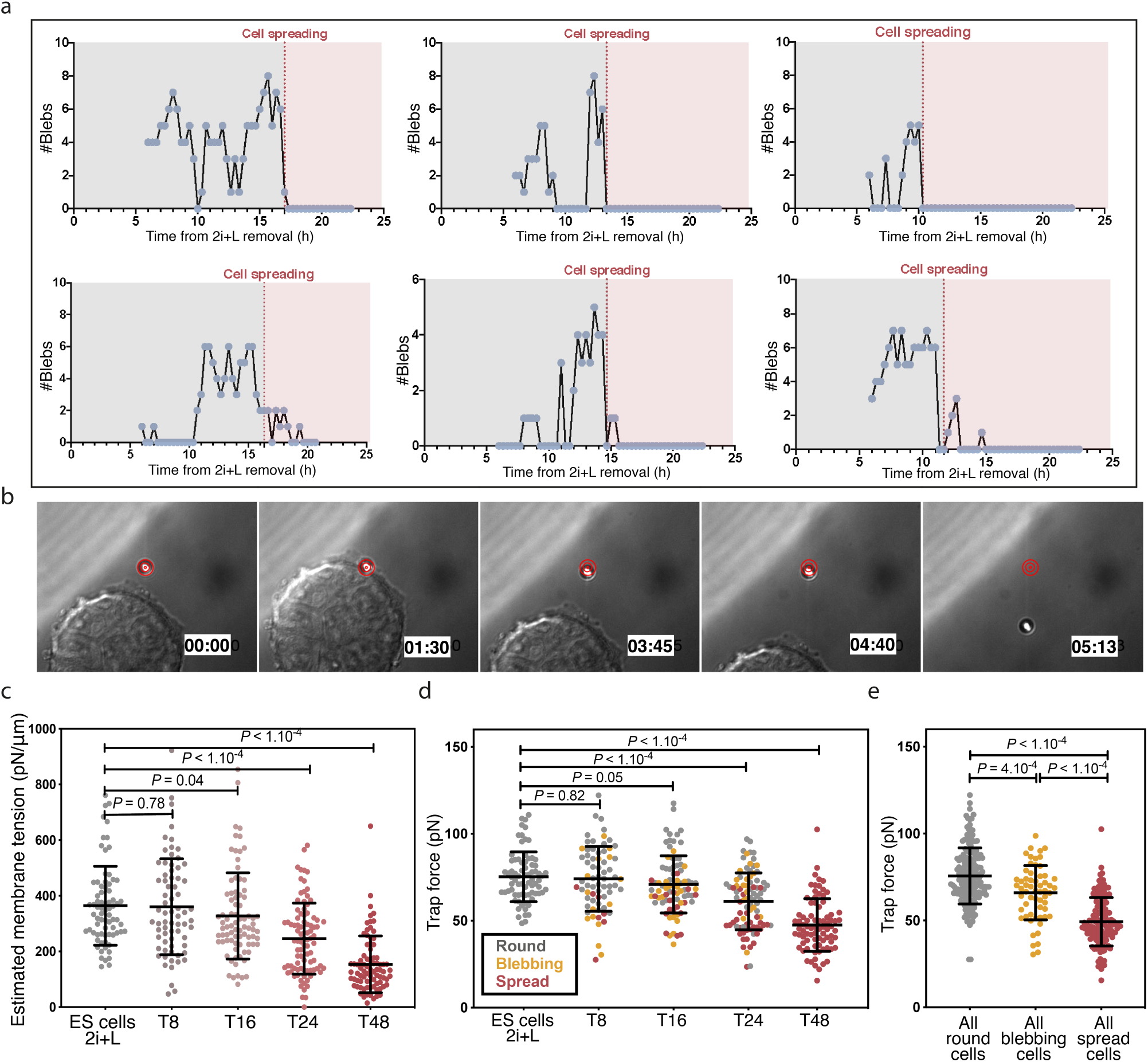
Membrane tension during exit from naïve pluripotency. **a**, Quantification of blebbing in six different ES cells expressing LifeAct-GFP during exiting naïve pluripotency. Red dotted line indicates approximate time of cell spreading. Bleb numbers per cell significantly decrease once the cell is spread (pink shaded area) **b**, Time lapse of a membrane tension measurement in a colony of ES cell. Time is in seconds. A red target is placed at the centre of the trap to help visualize bead displacement. **c**, Estimated membrane tension from the measurements presented in Fig. 1h. with the addition of T8 and T16. Membrane tension was estimated from the trap force measurements following equations described in (32). Arbitrary bending modulus of 20 pN/μm was picked in agreement with the literature (32) (37). (Means ± SD, N=3). **d**, trap force measurements of ES, T8. T16, T24 and T48 cells. Data are colour coded based on cell shape (grey = round, orange = blebbing, red = spread)(N=5). **e**, same data as in d panel but with all the data grouped by shape (e.g. all blebbing cells correspond to all the blebbing cells observed in T8, T16 and T24). *P* values are calculated by Welch’s unpaired student t-test and indicated in the figure. Scale bars = 10μm.

**Supp Fig. 3:**
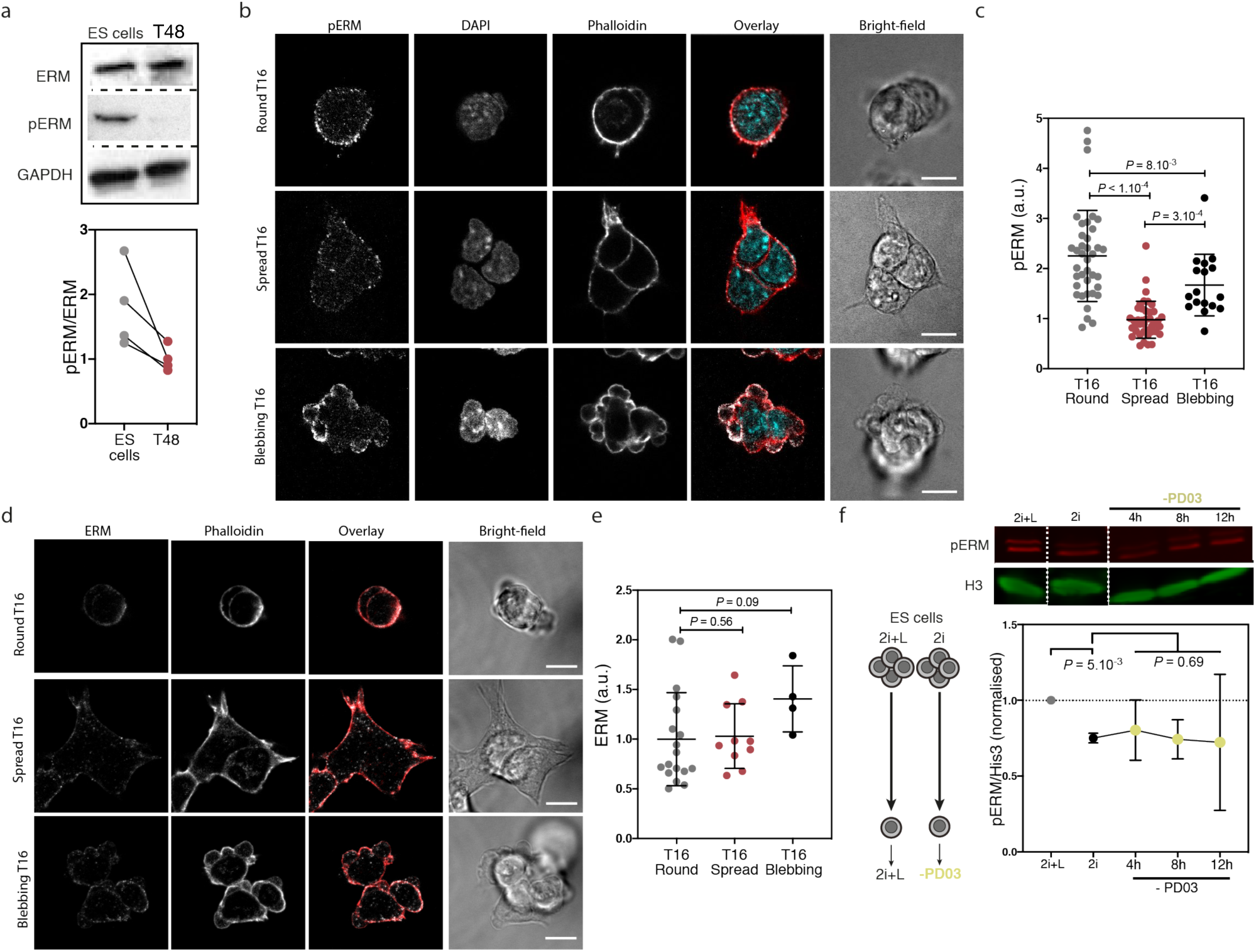
ERM is dephosphorylated during exit from naïve pluripotency. **a**, Representative western blot for ERM, pERM and GAPDH in ES cells and T24 cells and corresponding quantification (N=4, lines connect data points from individual experiments) **b**, Images of a single z-plane in fixed T16 cells. Three different cell morphologies are observed at T16: round, spread and blebbing. Cells were immunostained for pERM, and stained with DAPI and phalloidin. **c**, Estimated pERM and density in round, spread and blebbing T16 cells. Protein density was estimated from the mean intensity of sum z projections, corrected for background, and normalised to the mean of T16 round cells. (Means ± SD, N=2). *P* values are calculated by Welch’s unpaired student t-test and indicated in the figure. **d, e**, Similar as b and c, with ERM instead of pERM. **f**, Left, schematic of the experimental setup. Cells were equilibrated by passaging for two passages in 2i without LIF in order to ensure that LIF, a cytokine, was entirely washed out. Right, Fluorescent Western Blots and quantification for pERM and His3 of ES cells cultured in 2i and replated in N2B27 with Chir only. Data are normalised to cells cultured in 2i+L (N=4). **c**, Fluorescent Western Blots and quantification for pERM and His3 of ES cells cultured in 2i+L and replated in N2B27 with PD03. Data are normalised to cells cultured in 2i+L (N=4). Graphical data represents mean ± SD. *P* values are calculated using a two-way analysis of variance (ANOVA), using experiment repeats and media condition as variables, and indicated in the figure. Scale bars = 10μm.

**Supp Fig. 4:**
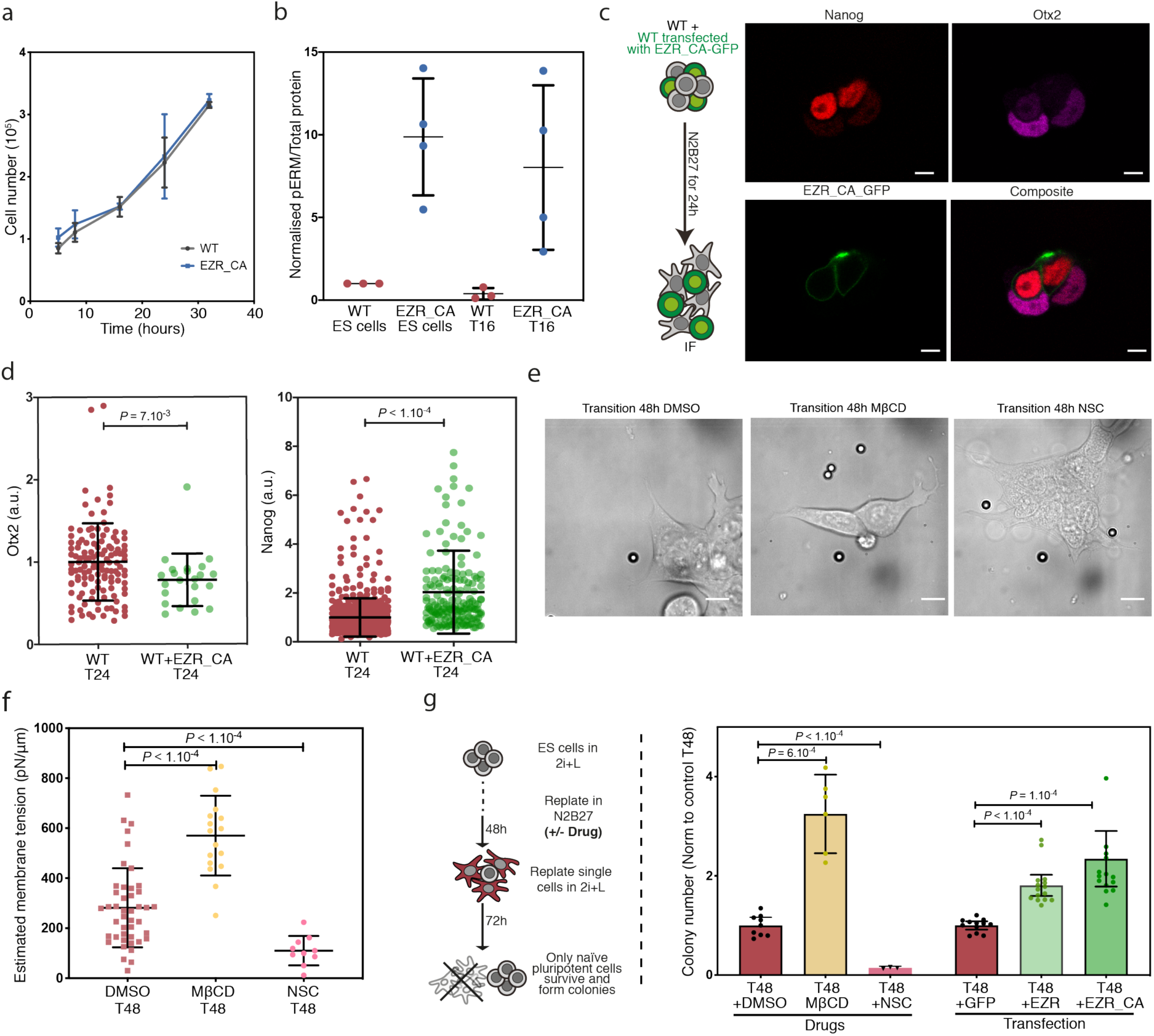
Increasing membrane tension prevents exit from naïve pluripotency. **a**, Proliferation curves of WT and iEZR_CA ES cells (N=3). **b**, Quantification of phospho-ERM normalised to total protein levels in WT and iEZR_CA ES and T16 cells. Data are normalised to the mean of WT ES cells (N=4). **c**, Representative single z-planes of a mix of control and EZR_CA transfected (positive for the EZR_CA _GFP) T24 cells immunostained for Otx2 and Nanog. **d**, Quantification of Nanog and Otx2 expression in T24 cells transfected or not with EZR_CA_GFP, normalised to T24 control mean (N=2). **e**, Representative images of T48 cells treated with DMSO, NSC23766 or MβCD. MβCD was used at 1 mM for 1 h every 24 h. NSC23766 was used at 25 μM for 1 h every 24 h (N=3). **f**, Membrane tension measurements of T48 cells treated with DMSO, NSC23766 or MβCD at similar concentrations as in e. Membrane tension was estimated as in Fig. S3B (N=2). **g**, Left, schematic of clonogenicity assay used to measure efficiency of exit from naïve pluripotency. Right, quantification of surviving replated cells in a clonogenicity assay using T48 cells and different treatments to affect membrane tension. T48 cells replated from N2B27+DMSO were used as control for drug treatment and T48 cells transfected with GFP were used as control for transfections. (N=3). Graphical data represents mean ± SD unless specified otherwise). Each condition is normalised to their respective control. *P* values are calculated by Welch’s unpaired student t-test and indicated in the figure. Scale bars = 10μm.

**Supp Fig. 5:**
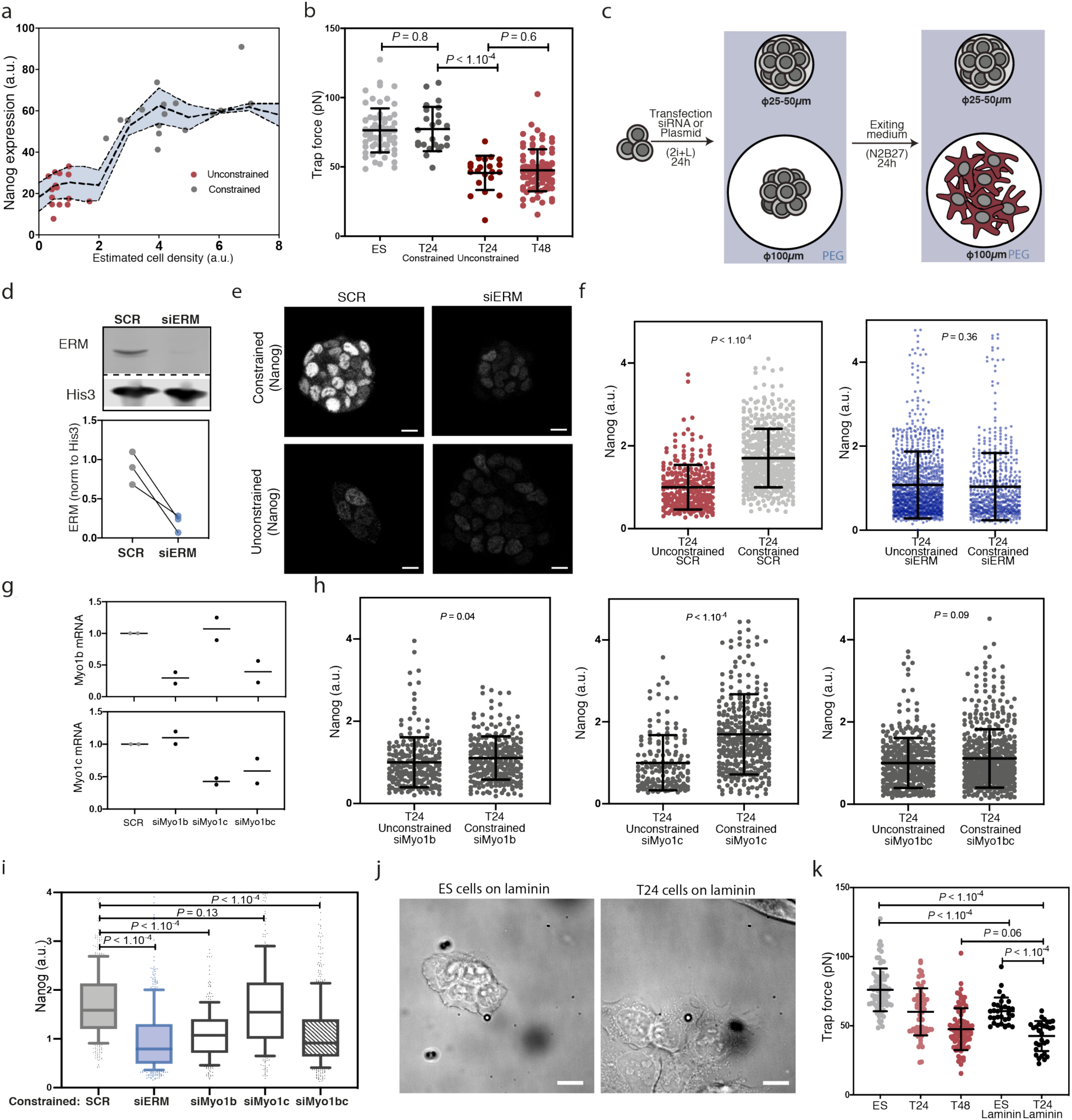
Maintaining a high membrane tension impairs exit from naïve pluripotency independently of cell shape. **a**, Same Nanog expression as in Fig. 2h plotted in function of estimated cell density. Cell density was estimated by counting the number of cells and dividing by the pattern area. Data are colored based on size of pattern (red for 100 μm diameter and grey for 25-50 μm diameter). **b**, Membrane tension measurements in ES cells and T24 cells either grown on micropatterns (small – constrained or large – unconstrained), or cultured in an open gelatin-coated dish (ES and T48). Data for cells on gelatine are from Fig. 1H. (N=2). **c**, Schematic of the micropatterning assay combined with transfection of plasmids or siRNA. **d**, Fluorescent Western blot (inverted contrast) for ERM and Histone 3 in cells transfected with siSCR and siERM showing the efficiency of the knockdown. **e**, Representative images of single z-planes in fixed T24 cells treated with either SCR or ERM siRNA and cultured on micropatterns. **f**, Quantification of Nanog expression in fixed T24 cells transfected with either SCR or siERM and cultured on micropatterns. Data are normalised to cells cultured on 100 μm diameter patterns (Unconstrained). Data for siERM are the same as in Fig. 2i. (Mann-Whitney U test was used, N=3). **g**, Quantification of Myo1b and Myo1c RNA by qPCR assay in ES cells treated with either SCR, siMyo1b, siMyo1c and siMyo1b+siMyo1c. Data were normalised to Actb expression. **h**, Similar to f with T24 cells either transfected with siMyo1b, siMyo1c or siMyo1b+siMyo1c (N=2). (I) Boxes and whiskers with 10-90 percentile plot of previous data. Only data for cells constrained on small micropatterns are shown (Mann-Whitney U test). **j**, Representative images of ES and T24 cells cultured on laminin. **k**, Trap force measurements in ES cells and T24 cells cultured on either gelatine or laminin. Data for cells cultured on gelatine are the same as Fig. 1H. (N=3). Graphical data represents mean ± SD unless specified otherwise. *P* values are calculated by Welch’s unpaired student t-test (unless specified otherwise) and indicated in the figure. Scale bars = 10μm.

**Supp Fig. 6:**
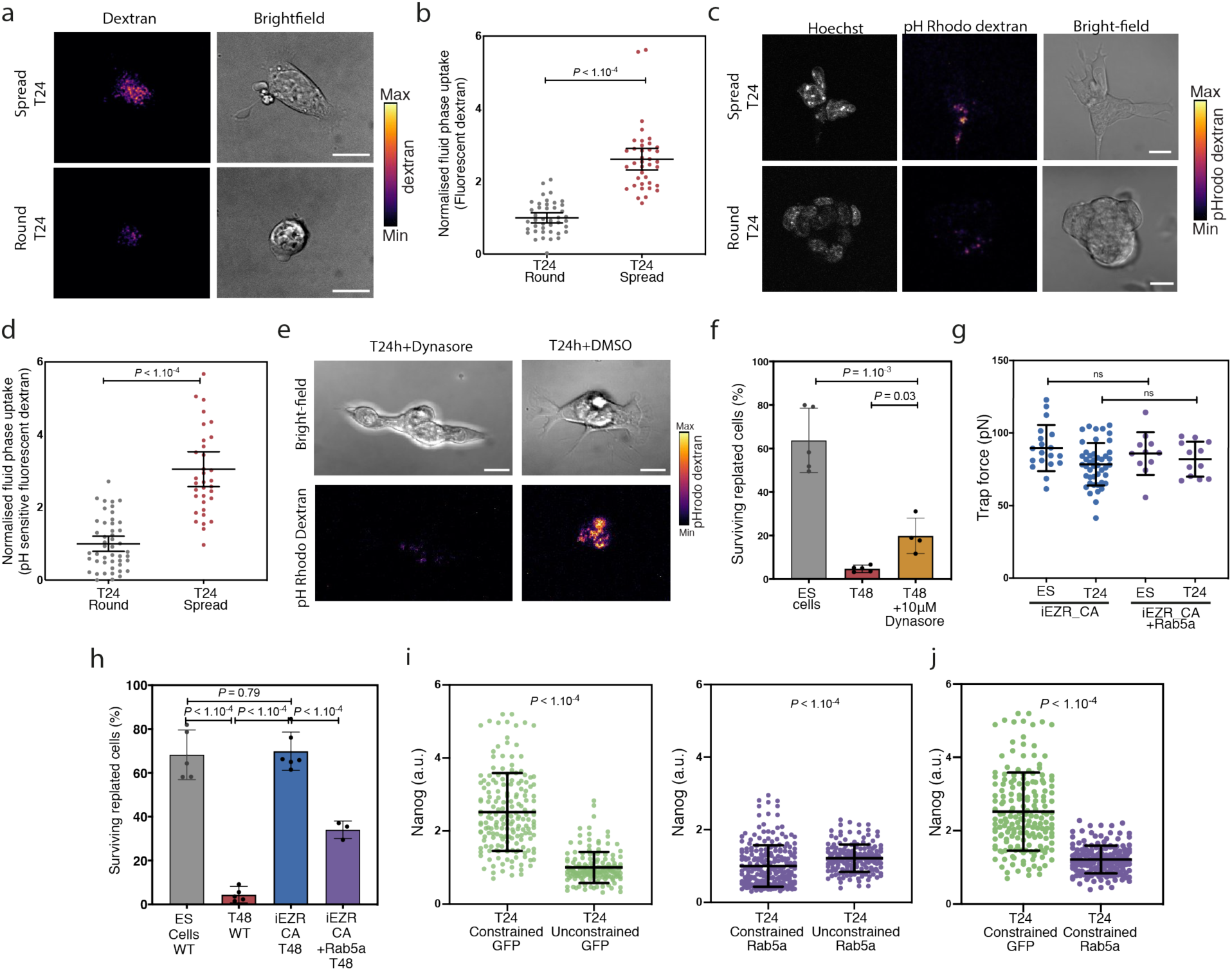
Perturbing endocytosis results in defects in exit from naïve pluripotency. Increasing endocytosis levels with Rab5a can rescue defects of exit in the EZR_CA cells and of cells physically constrained on micropatterns. **a**, Representative images of sum z-projections in round and spread T24 cells during a liquid phase uptake assay using fluorescent dextran. **b**, Quantification of liquid phase uptake using fluorescent dextran in T24 cells. Data are separated based on cell shape (round or spread). (Mean ± 95% Confidence Interval, N=2). **c**, Similar as a, but using pH sensitive fluorescent dextran (pH Rhodo) **d**, Similar as b, but using pH-sensitive fluorescent dextran (N=2). **e**, Representative images of sum z-projections in representative T24 cells treated with either DMSO (control) or with 10 μM of dynasore. **f**, Quantification of the percentage of surviving replated cells in a clonogenicity assay using ES cells and T48 cells treated or not with 10 μM concentrations of dynasore. Cells treated with dynasore for more than 24h started showing significant growth defects which might explain the low colony numbers observed. Data are normalised to non-treated T48. (N=4). **g**, Trap force measurements of iEZR_CA cells transfected with Rab5a in 2i+L and at T24. Data for WT and iEZR_CA cells are from Fig. 2B (N=2). **h**, Quantification of the percentage of surviving replated cells in a clonogenicity assay using ES and T48 cells, WT or iEZR_CA. transfected or not with Rab5a. Data for iEZR_CA T48 cells are from Fig. 2E. (N=3). **i**, Quantification of Nanog expression of fixed T24 cells transfected with GFP as control or Rab5a, cultured on micropatterns. This shows that Rab5a transfection can also rescue exit defects in constrained cells cultured on micropatterns. Data are normalised to cells cultured on 100 μm diameter patterns (Unconstrained) (Mann-Whitney U test is used; N=3). **j**, Box and whiskers with 10-90 percentile plot of data in panel i. Only data for cells “constrained” on small micropatterns are shown (Mann-Whitney U test). Graphical data represents mean ± SD unless specified otherwise. *P* values are calculated by Welch’s unpaired student t-test (unless specified otherwise) and are indicated in the figure. Scale bars = 10μm.

**Supp Fig. 7:**
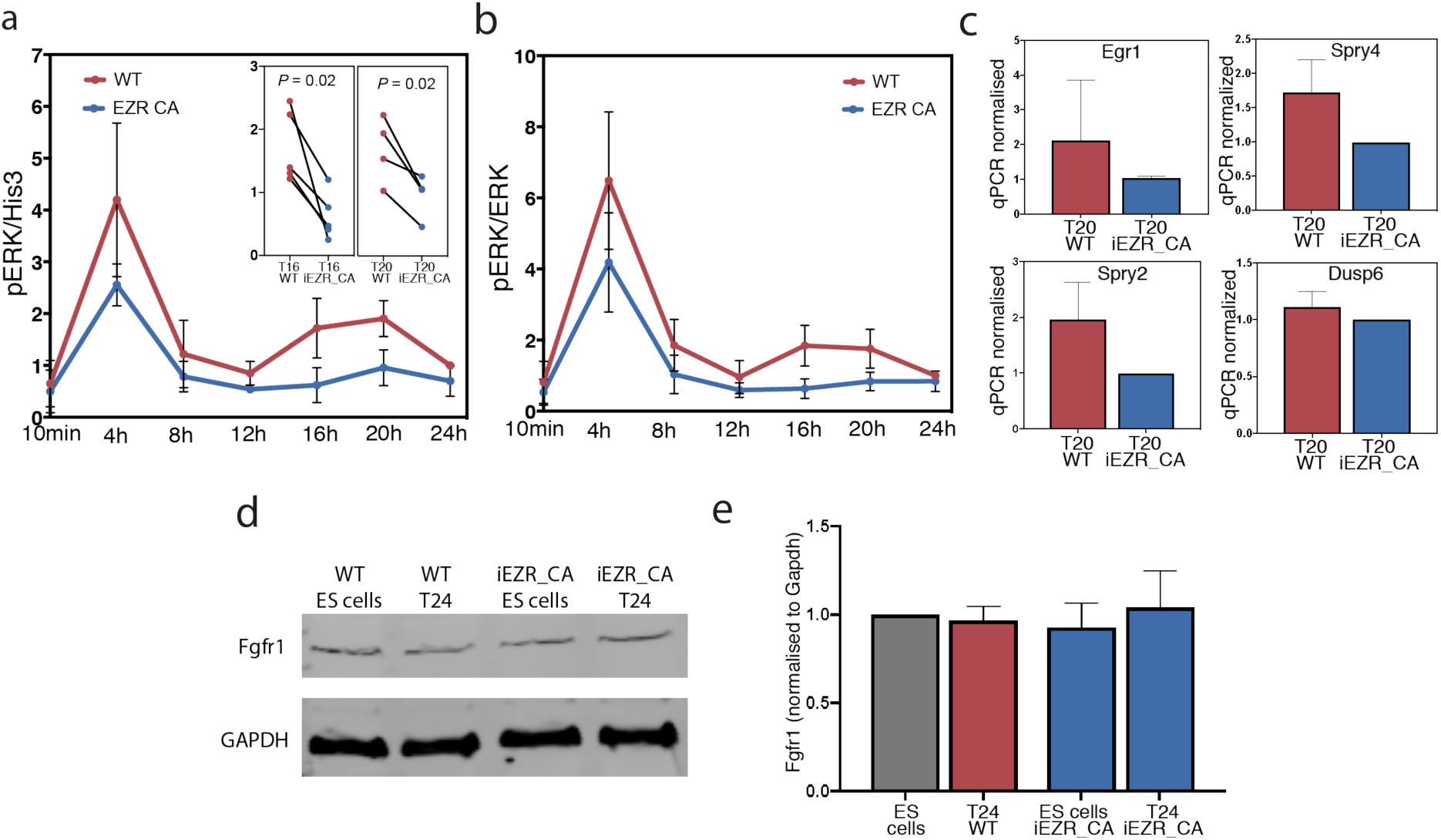
ERK signalling and FGFR1 expression in exit from naïve pluripotency. **a**, Quantification of phospho-ERK normalised to Histone 3 levels across several fluorescent Western blots (including Fig. 4a) done on WT and iEZR_CA cells at different timepoints during exit from naïve pluripotency. The first peak of ERK activation is likely a direct effect of the removal of the inhibitors. The timing of the second peak approximately coincides with the timing of the decrease in membrane tension and cell spreading (N=5). Inset, data for T16 and T20 cells where values for WT and iEZR_CA in individual experiments are connected by a line (N=5). **b**, Similar as panel a, for phospho-ERK normalised to total ERK levels. **c**, Quantification of mRNA expression for Egr1, Spry4, Spry2 and Dusp6 measured by qPCR in WT and iEZR_CA T20 cells. Expression was normalised to Actb and data are normalised to the mean for iEZR_CA T20 cells (N=3). **d, e**, Fluorescent Western blot (inverted contrast) for Fgfr1 and GAPDH in WT and iEZR_CA ES and T24 cells, and corresponding quantification (right, data are normalised to GAPDH). (N=3). *P* values are calculated by Welch’s unpaired student t-test (unless specified otherwise) and indicated in the figure. Graphical data represents mean ± SD unless specified otherwise.

**Supp Fig. 8:**
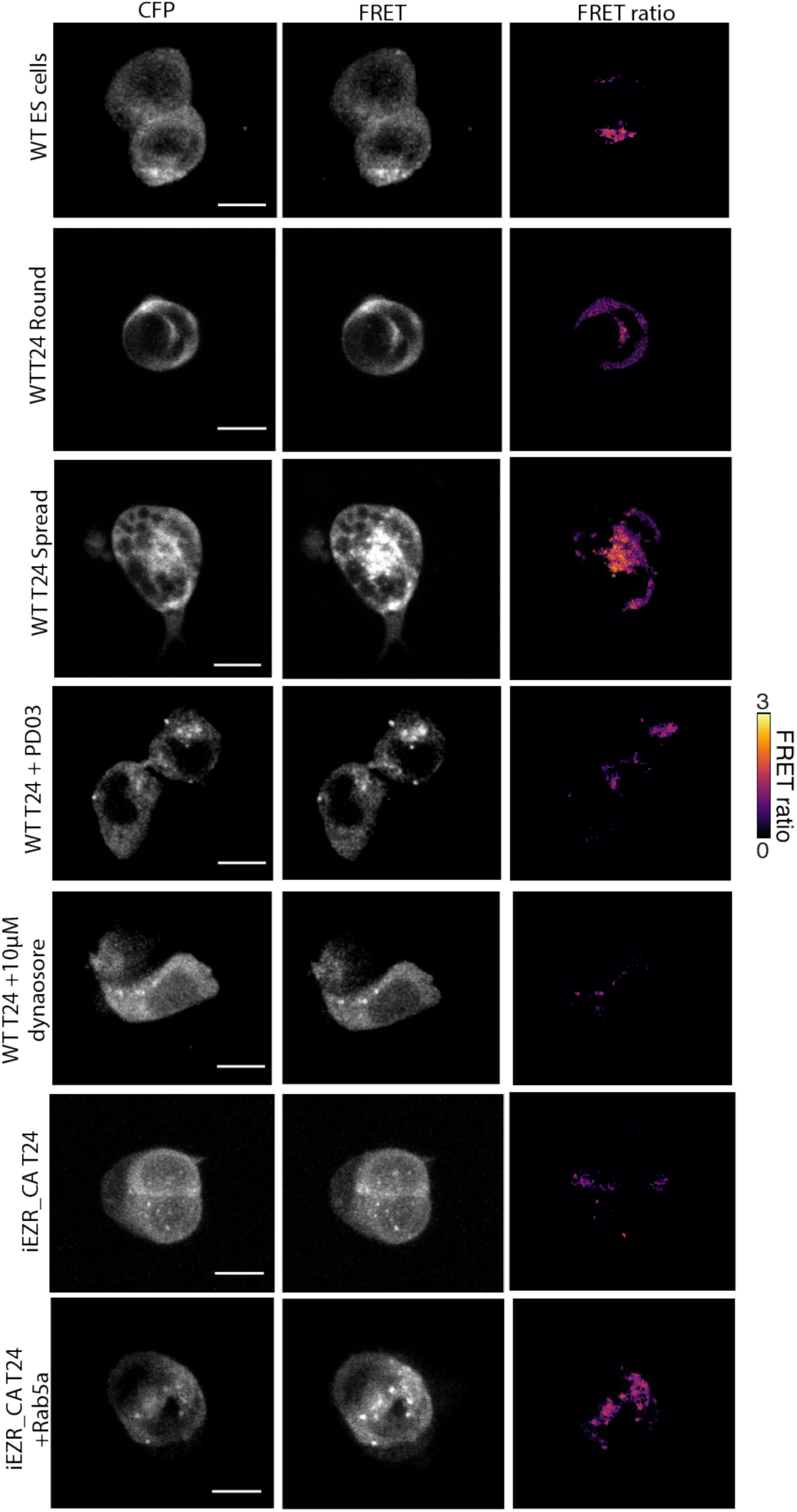
ERK activation at early endosomes. Representative images of z projections of fixed ES cells in different conditions. PD03 was used at 1 μM. Dynasore was used at 10 μM. Displayed pictures have been smoothed (Gaussian blur). Scale bars = 10 μm.

**Supp Fig. 9:**
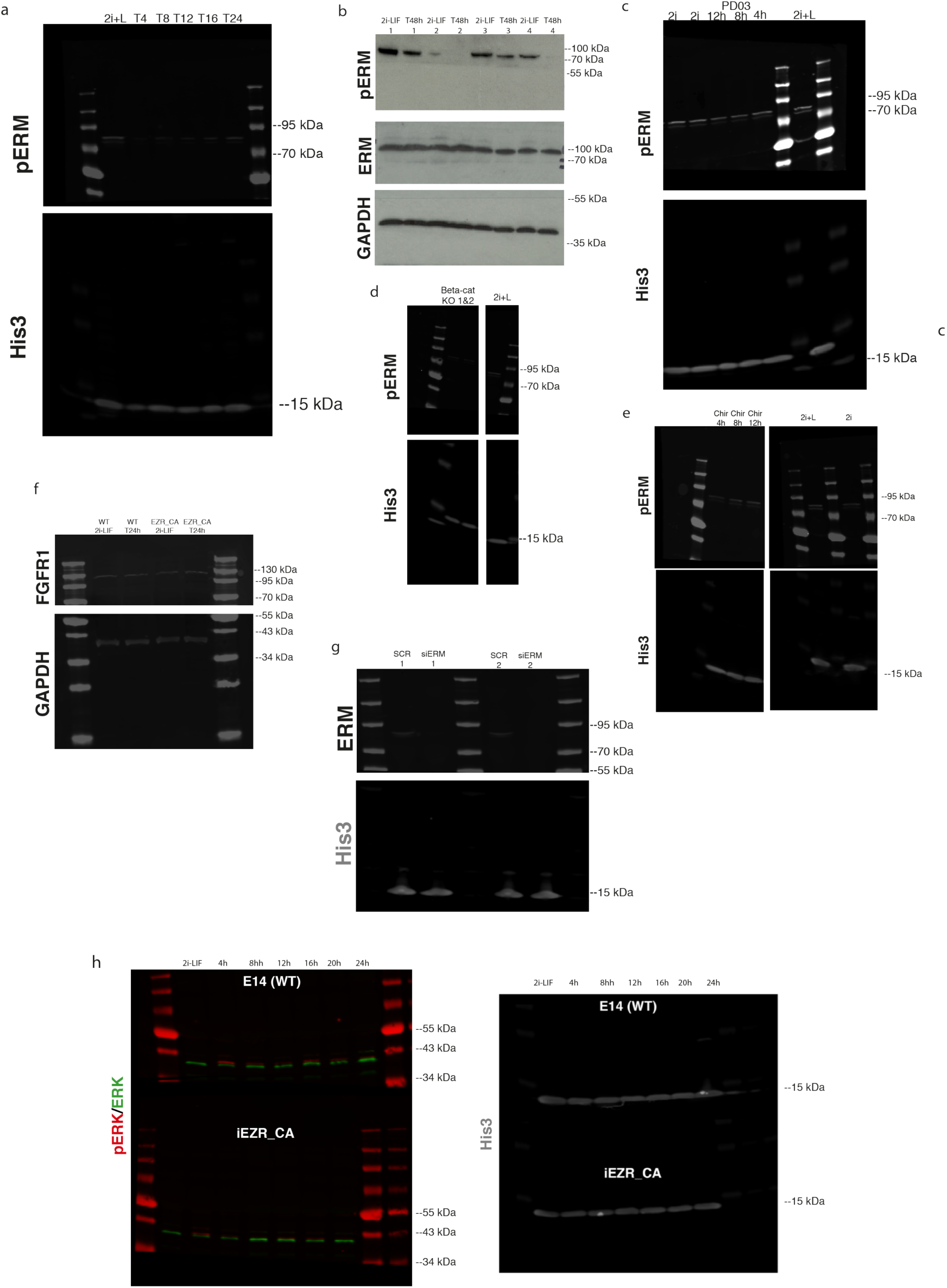
Uncropped Western Blots. Uncropped Western Blots corresponding to: **a**, Fig. 1h. **b**, Sup Fig. 3a. **c**, Fig. 1i. **d**, Fig. 1j. **e**, Sup Fig. 3f. **f**, Sup Fig. 7d. **g**, Sup Fig. 4b. **h**, Fig. 4a.

